# Model supports asymmetric regulation across the intercellular junction for collective cell polarization

**DOI:** 10.1101/2024.06.03.597091

**Authors:** Katherine Levandosky, Calina Copos

**Affiliations:** Department of Mathematics, Northeastern University, Boston, MA, USA; Department of Biology, Northeastern University, Boston, MA, USA

## Abstract

Symmetry breaking, which is ubiquitous in biological cells, functionally enables directed cell movement and organized embryogenesis. Prior to movement, cells break symmetry to form a well-defined cell front and rear in a process called polarization. In developing and regenerating tissues, collective cell movement requires the coordination of the polarity of the migration machineries of neighboring cells. Though several works shed light on the molecular basis of polarity, fewer studies have focused on the regulation across the cell-cell junction required for collective polarization, thus limiting our ability to connect subcellular interactions to tissue-level dynamics. Here, we investigate how polarity signals are communicated from one cell to its neighbor to ensure coordinated front-to-rear symmetry breaking with the same orientation across the group. In a theoretical setting, we exhaustively search a variety of intercellular interactions and identify the conditions required for the Rho GTPase signaling module and/or cytoskeletal dynamics to achieve either co-alignment arrangement or supracellular arrangement of the polarity axes in a group of 2 and 4 cells. Our work shows that only asymmetric regulations are favorable – such interactions involve either up-regulation of the kinetic rate of complementary polarity components or opposite regulation of the kinetic rates of the same polarity components across the cell-cell junction. Surprisingly, our results hold if we further assume the presence of an external stimulus, intrinsic cellular variability, or larger group size. The results underline the potential of using quantitative models to probe the molecular interactions required for macroscopic biological phenomena. Lastly, we posit that asymmetric regulation is achieved through junction proteins and predict that in the absence of cytoplasmic tails of such linker proteins, the likeliness of doublet co-polarity is greatly diminished.

**Author summary:** Cells of the developing embryo undergo a highly complex chain of events that define their correct shape and positioning. Among these events, a crucial role belongs to coordinated cell movement of cells of different lineages over short and long distances to give rise to mature organs and organ systems. During collective movement, individual cells typically engage their autonomous polarity machinery, while being connected to their neighbors through adhesive cell-cell interactions. Despite advances in revealing the cell-cell interactions required for collective cell migration, a comprehensive picture of the molecular basis of intercellular communication for collective guidance is missing. To address this question, we devise a generalized mechanochemical model for cell polarity in a doublet and investigate how polarity signals are transmitted from one cell to another across seemingly symmetrical junctions. We have chosen to screen through all possible intercellular conditions of the Rho GTPase signaling circuit and/or cytoskeletal dynamics. Our systematic approach provides information on over 300 distinct conditions and reveals the intercellular regulation provided by junction proteins. In addition to predicting that only asymmetric interactions favor co-polarization, ensuring movement of the group in the same direction, our analysis also highlights the need for additional regulatory mechanisms for larger cell groups.

## Introduction

From bacterial to mammalian cells, cell polarity is essential in a multitude of functional contexts, including cell migration, division and differentiation, and development [1–5]. Cell polarity is manifested in molecular and morphological asymmetries across the cell [6, 7]. One fundamental question related to cell polarity is how an initially symmetric cell can spontaneously establish a polarized state, with a well-defined cell front and rear, but also show sensitivity to external guiding cues [8]. Cells are also known to engage in collective migration, which necessarily requires coordination of the individual cell’s direction of movement by signaling across symmetric cell-cell junctions. Previous studies have shown that vectorial signaling requires mechanical coupling between cells through cadherin dependent cell-cell junctions [9–12]. This raises a second fundamental question: What are the underlying biochemical and/or mechanical interactions at cell-cell junctions that support co-orientation of polarity axes such that all cells in a group initiate migration in the same direction?

The first question is well studied, both on the conceptual level with theoretical approaches, and on the experimental level, by characterizing signaling pathways. The polarization of an initially non-polarized cell is a symmetry breaking phenomenon: in the case of essentially isotropic cells, the continuous angular symmetry is broken by polarization, which can happen spontaneously [8, 13], but is often controlled by upstream guiding cues [14], and noise can play an important role [15]. Polarity establishment arises primarily through the localization of specific proteins and lipids in the cell to specific regions of the plasma membrane, and often precedes motility [6, 7]. While the detailed molecular mechanisms differ between organisms, they involved a relatively small, conserved set of proteins – here, we focus on the Rho molecular circuit [3, 16] and specifically the GTP-GDP cycling of small GTPases Rac1, which promotes lamellipodial protrusions at the migrating front, and RhoA, which promotes contractility at the rear, (Fig. 1a) – these proteins will be referred to as Rac and Rho, respectively, henceforth. However, cell polarization can also be associated with the rearrangement of the actin cytoskeleton, in which branched actin filaments form at the cell front while actomyosin contractile bundles segregate to the cell rear [4, 17, 18] (Fig. 1a). Just as diffusible chemical activators and inhibitors trigger biochemical instabilities, mechanical instabilities can arise due to stochastic fluctuations in actin filament densities or mechanical feedback between motor proteins and cytoskeleton elements [19]. In mechanically driven polarity systems, cells polarize due to the mechanical forces and the actin flow generated by these forces [4, 20, 21]. Two classic cases involving cytoskeleton-driven polarization are the formation of actin comet tails by intracellular pathogens [22] and the directional locomotion of keratocytes [4, 13, 23]. In both cases, the mechanical properties of the actin cytoskeletal network appear sufficient for polarization, which can be triggered by stochastic or induced asymmetries in the mechanical network. Although cell polarity can emerge from systems that are either chemical or mechanical, in many cases cell polarity depends on the interplay between the two to robustly break symmetry to initiate locomotion [24–28].

**Fig 1.**
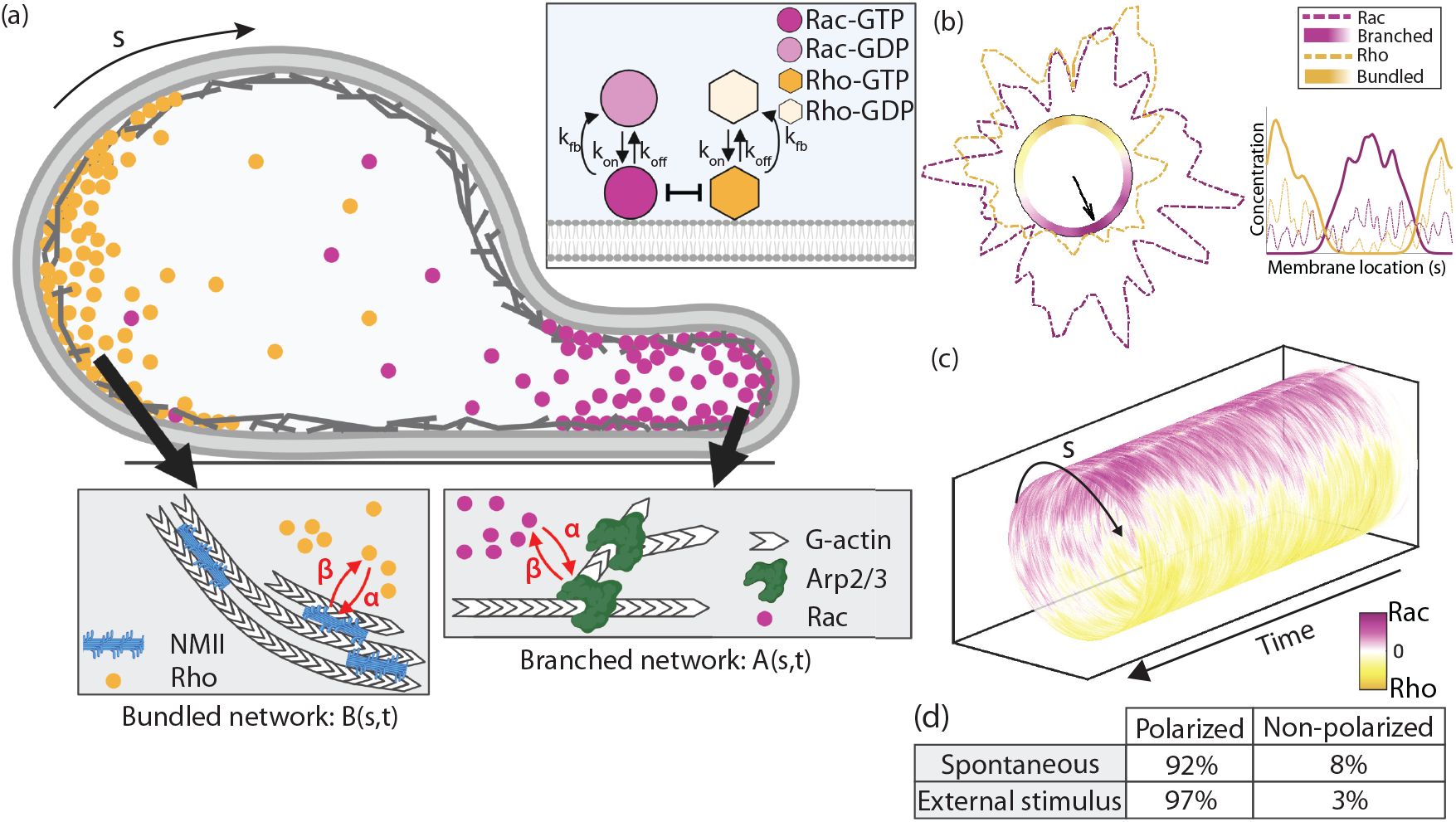
Spontaneous polarity in individual cells. (a) Schematic representation of front-to-rear polarity markers. The abundance of two polarity protein markers at the two distinct sites is shown with Rho (yellow) and Rac (magenta). Local, bidirectional crosstalk between Rho and actomyosin bundles and Rac and Arp2/3 branched F-actin networks (red arrows) ensures the simultaneous spatial organization of two distinct F-actin networks supporting the formation of a cell front and rear. The inset shows the GTP-GDP cycling dynamics of Rho GTPases in the model. (b) The outcome of one model realization. The cell membrane is shown in black. The branched and bundled F-actin networks are plotted inside the cell, while the GTPase concentrations are shown outside the cell membrane (dashed). The front-to-rear polarity axis is drawn from the cell center to the middle point of the region where the branched F-actin network is above threshold level (black arrow). The inset shows concentrations of all 4 polarity markers along the cell membrane. (c) Rac and Rho concentrations in space and time averaged over 100 realizations – a cylinder slice corresponds to the Rho GTPase concentrations at one fixed time point along the plasma membrane. (d) Summary of polarization probability for 100 realizations of the model.

In collective migration, each cell individually contributes to the group’s migration by first breaking symmetry and establishing a polarity axis while maintaining physical contact with neighboring cells. For the group to move together in the same direction, further mechanisms are required for coordination of the polarity of their autonomous migratory machineries. Experimental work has focused on uncovering the links between cell signaling pathways and collective cell movement. In epithelial layer sheets, ERK signaling waves are tightly connected to mechanical forces to ensure collective migration [29]. In collectively migrating human umbilical vein endothelial cells, physical membrane protrusions termed “cadherin fingers” interconnect the rear of leading cells to the front of follower cells [30]. These VE-cadherin rich structures are deeply connected to the actin cytoskeleton of both follower and leader cells and are thought to regulate Arp2/3 actin polymerization. Another clue into the intercellular coordination of the polarity pathway for collective migration comes from chemotaxing neural crest cells [31]. As neural crest cells “chase” placodal cells, before cell-cell contact, neural crest cells have high, localized Rac activity at the cell front but after contact, junction proteins (N-cadherins) inhibit Rac localization. Importantly, in cell “trains”, as exemplified by the migration of neural crest cells, collision and contact inhibition of locomotion (CIL) have been demonstrated to play a role *in vivo* by maintaining coherent directional migration of groups of cells [32]. A number of theoretical models have been developed to study the emergence of directed collective migration, reviewed in [33]. In particular, one model has focused on identifying the mechanisms, chemical and/or mechanical, that can account for CIL in interacting cell groups in confinement [34]. Despite these combined efforts, the driving mechanisms to ensure coordination of collective symmetry breaking prior to migration remain elusive. In Table 1, we review a subset of the experimentally postulated intercellular signaling interactions between neighboring cells for collective migration.

**Table 1.**
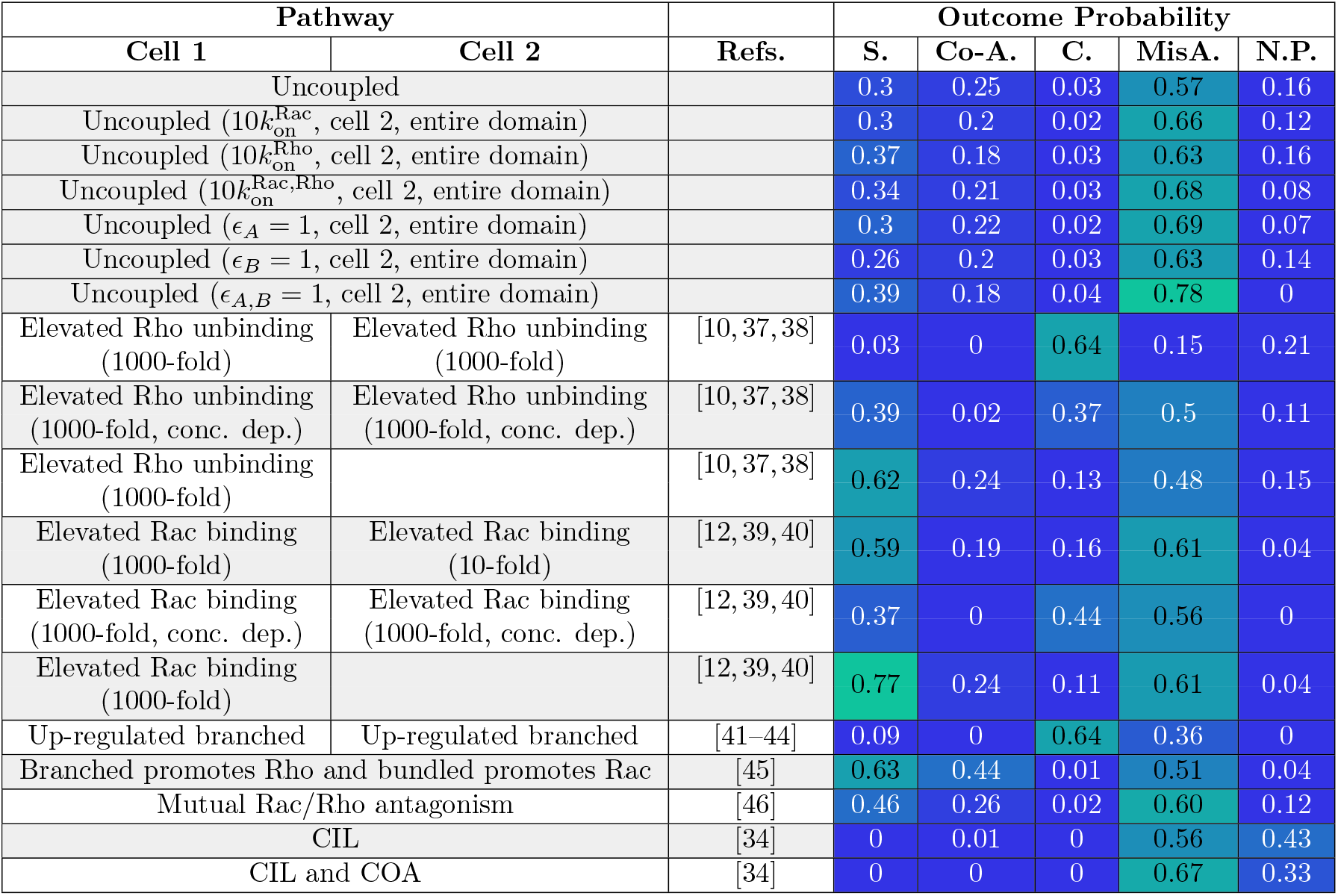
A subset of molecular-based pathways of cell-cell coupling for a pair of cells are listed based on experimental findings. The speculated couplings are implemented in the model and the outcome probabilities of supracellular (S.), co-alignment (Co-A.), collision (C.), misalignment (MisA.), or non-polarized (N.P.) arrangement for 100 realizations of the model are presented.

Here, rather than cells being pulled or pushed along, we exhaustively search the intercellular biochemical and/or mechanical conditions for neighboring cells to coordinate their symmetry breaking processes ahead of movement. Specifically, we identify the simplest conditions at the cell-cell junction that ensure individual polarity axes are co-aligned towards a common direction across the cell group. We use a previously developed mechanochemical model for polarization of an individual cell [35] (Fig. 1b-d). The model is extended to a pair of cells and an exhaustive search of the interactions at the cell-cell junction are evaluated, including interactions which rely on the biochemical circuit, the mechanical circuit, or both. Our results identify a small set of asymmetric interactions of the polarity markers which favor co-alignment or supracellular arrangement of front-to-rear axes in the doublet. Surprisingly, our finding holds even if we assume intrinsic variability between the cells or an external signal orienting polarization rather than spontaneous polarization. We posit that these types of intercellular couplings at symmetric cell-cell junctions arise from mechanical forces at the cell-cell junction, which asymmetrically regulate the Rho signaling pathway in neighboring cells. In addition, we use our model to study collective polarization for larger groups of cells. One would expect that groups of 4 cells behave similarly to doublets, but, surprisingly, initial geometric arrangement also plays an important role. We find that groups of 4 cells in a square (over linear) arrangement exhibit a wider variety of behaviors, ranging from co-alignment to clockwise or counterclockwise rotational alignment. This can be understood as due to the larger number of degrees of freedom, almost identical to behavior of cells in confined environments rather than flat surfaces [36]. Our findings suggest that additional regulatory mechanisms, perhaps CIL, are at play to sustain co-alignment organization of polarity axes in tissues.

## Model

### Molecular ingredients of the mechanochemical polarization model

The individual cell polarization mechanism is based on a mechanochemical model [35]. It assumes a biochemical signaling circuit, based on small GTPase active-inactive cycling, with positive, local, bidirectional feedback into a mechanical circuit, based on “frontness”/”backness” cytoskeletal dynamics (Fig. 1a). The quantitative mechanism suggested by the model is simple: branched (bundled) actin networks support recruitment of Rac (Rho) molecules to the membrane, so Rac (Rho) molecules tend to segregate into separate parts of the cell. In turn, neither network can invade the other’s spatial domain, because Rac (Rho) molecules engage the branched (bundled) network. These dynamics lead to robust spontaneous polarization in an individual cell (Fig. 1b,c) as well as in the presence of an external stimulus (Fig. 1d). Briefly, we outline the dynamics assumed in each sub-circuit of the mechanochemical polarization model.

a. <u>Rho GTPase signaling circuit</u>: The biochemical signaling dynamics are based on the well-studied GTP-GDP cycling of the small GTPases Rac and Rho. Each GTPase molecule cycles between two states: an active GTP-bound form, bound to the plasma membrane, and an inactive GDP-bound form, freely diffusing in the cytosol. Once the active GTP-bound molecule is bound to the membrane, it induces a positive feedback activation through recruitment of inactive molecules to the locations of membrane-bound active molecules. To capture the kinetics, we use a stochastic formulation to track the position (s) and the location of the activated, membrane-associated Rho GTPases at a given time *t*, 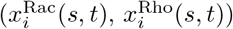 for the signaling molecule of index *i*. For the cell geometry, we consider a static circular one-dimensional periodic domain which represents the plasma membrane and a thin volume of cytoplasm adjacent to the membrane. The location of the molecules along the circle is given by *s*, the arc length. For each membrane-bound molecule *i*, we compute the time to its next reaction, which is exponentially distributed with rate:

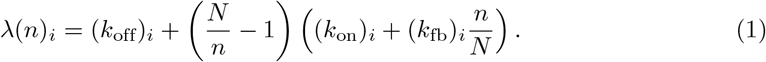

The rate *λ* should be interpreted as a reaction rate per membrane-bound molecule – hence, the unbinding rate per molecule is a constant *k*_off_, while the binding rate is proportional to the remaining fraction of available binding spots on the cell membrane with a constant of proportionality *k*_on_ + *k*_fb_*n/N* . The spontaneous association rate per molecule is constant, but the enhanced association per molecule due to its implicit dynamics of enhanced recruitment is proportional to the fraction of molecules that are still in the cytosol. The kinetic assumptions are also well justified by biological data and their rates are taken from literature. Between reaction events, the number of membrane-bound Rac (or Rho) molecules is constant, and the molecules diffuse freely on the membrane. A steric repulsion is enforced between Rac and Rho polarity molecules so that the two chemicals cannot cross paths at any moment in time. This assumption is also well justified by biological data [47–50]. To minimize assumptions, the reaction rates for both GTPases are identical. Parameter and implementation details are specified in Appendix A1 and in [35].
b. <u>Cytoskeletal dynamics circuit</u>: For the mechanical circuit, we describe the re-arrangement of the F-actin network as coupled reaction-diffusion equations, which describe the densities of branched protrusive actin network, *A*(*s, t*), and contractile bundled actomyosin network, *B*(*s, t*),

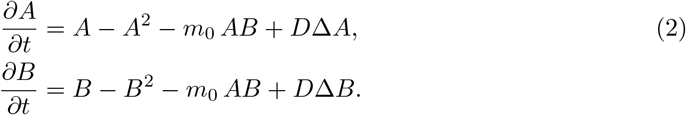

The branched network is assumed to be protrusive and devoid of myosin II motors, while the bundled network contains contractile actomyosin bundles that generate contractile forces and retract the cell’s posterior. Simply, their dynamics state that the rate of growth is proportional to the network concentration but limited due to finite molecular resources (e.g. branching complexes, myosin II motors, etc.) [51]. The model assumes that the competition terms, proportional to the product *AB*, stem from either mechanical competition or competition for limited molecular resources. Parameter *m*_0_ is the magnitude of competition between the networks. The last term accounts for the diffusion of the networks due to sliding and shuffling of bundled filaments in the contractile actomyosin network, as well as the random lateral displacements of the growing ends of the barbed filaments along the cell membrane [13].

The coupling between the biochemical signaling and mechanical circuits is achieved through a local positive feedback loop between Rac and branched actin (*x*^Rac^ and *A*), and Rho and bundled actin (*x*^Rho^ and *B*) (red arrows, Fig. 1a). The binding rates of small GTPases are not fixed but rather depend on the local concentration of each respective actin network. Similarly, the growth rate of each actin network is proportional to the local densities of membrane-bound polarity proteins. Without any further assumptions, it is easy to understand why polarity emerges. The cycling from active to inactive state leads to multiple activity sites [52]. The antagonism between Rac and Rho will lead to the formation of multiple distinct sites of active, membrane-bound Rac or Rho molecules. Simultaneously, branched, and bundled actin structures will compete to form and segregate along the cell boundary. Their dynamics are “winner take all”, and for stable co-existence of both structures further assumptions or nonlinearities are needed [13, 35, 53–55]. Here, their stability is improved by localization of Rac/Rho, respectively, which in turn leads to the collapse of multiple activity sites into a single Rac “frontness” patch and Rho “backness” patch in the opposite spatial location.

## Results

### The absence of intercellular interactions produces sporadic co-alignment of polarity axes in group

To probe the effect of junction protein complexes on regulating Rho signaling and/or F-actin network assembly at the cell-cell junction, we consider two cells that are equivalent: same biochemical kinetic rates and actin network parameters. The cells maintain static cell-to-cell contact and the intercellular junction is fixed to be 25% of the perimeter of each plasma membrane (Fig. 2a). In 100 simulated pairs (Fig. 2b), for each cell in the pair a front-to-rear polarity axis is determined from the cell center to the point on the plasma membrane that corresponds to the middle point of the region where the branched F-actin network is above threshold level (Fig. 2c, *C*_crit_ = 1, 10% of steady state concentration). To determine if the pair co-aligned their polarity axes, the orientation and angle difference between polarity axes are calculated. The possible outcomes are classified into four distinct non-overlapping scenarios (Fig. 2d):

**Fig 2.**
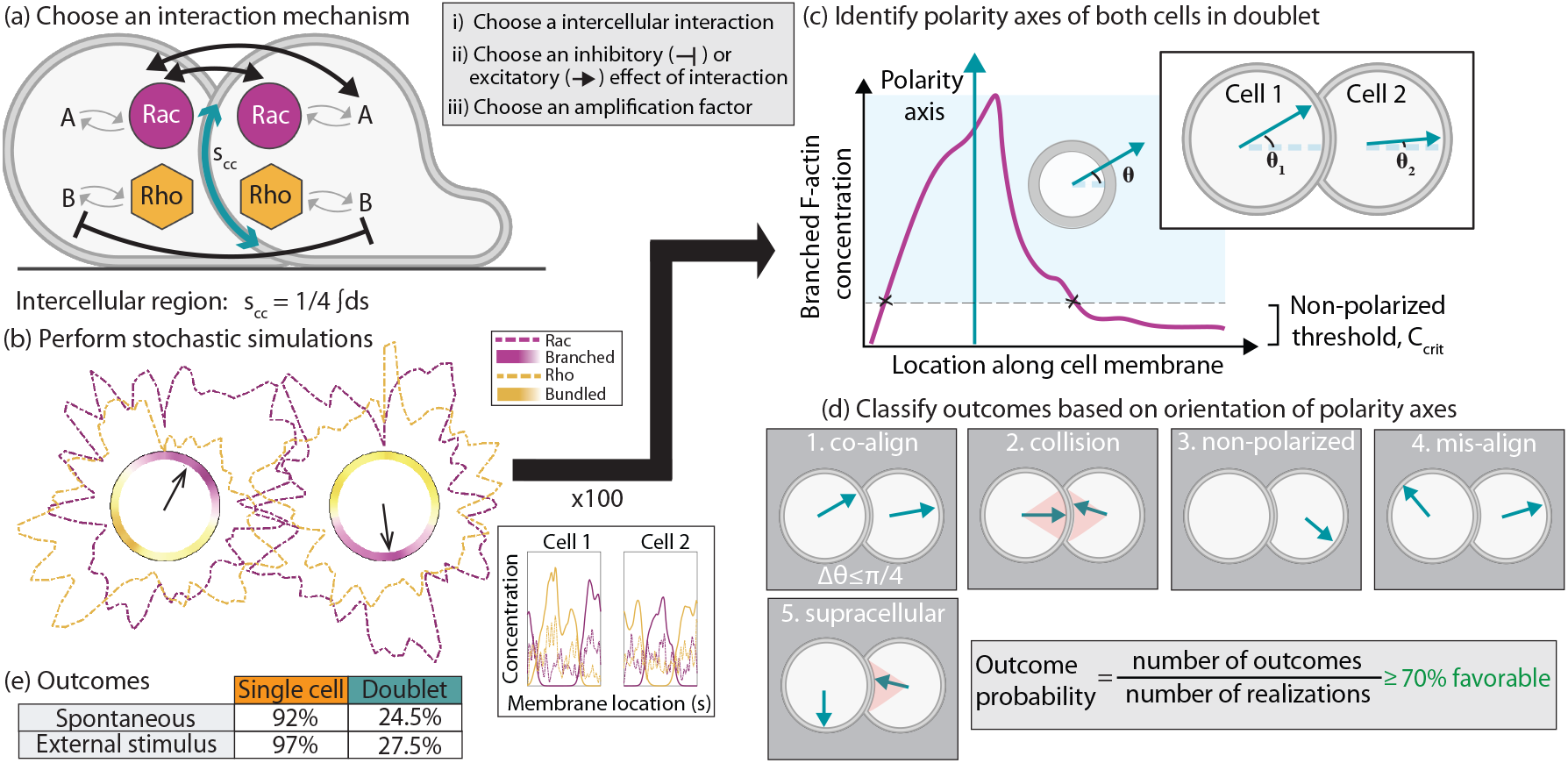
Schematic representation of workflow in a cell doublet. (a) A possible intercellular interaction in a pair of cells is selected. The interaction can be based on binding/unbinding of GTPases to the plasma membrane, assembly of F-actin networks, or a combination of the two. The interaction occurs in an intercellular region defined to be 25% of the circumference of either one of the cells. (b) Example of a simulation resulting in misalignment configuration. The cell membrane is shown in black. The branched and bundled F-actin networks are plotted inside the cell, while the GTPase concentrations are shown outside the cell membrane (dashed). The black arrows mark the polarity axes. The inset shows the concentration of all 4 polarity markers in both cells. (c) An outcome probability is calculated from 100 simulations. For each simulation, a front-to-rear axis is drawn from the cell center to the middle point of the region where the branched F-actin network is above a critical threshold value *C*_crit_. The angle opening from the horizontal to the polarity axis is calculated for both cells. (d) Based on the orientation of the polarity axes in both cells, an outcome is assignment. Possible outcomes are (1) co-alignment, (2) collision, (3) non-polarized, (4) misalignment, and (5) supracellular arrangement. (e) Summary of probability outcome for single cells (number of polarized cells/number of realizations) and doublets (number of doublets with both cells polarized in co-alignment arrangement/number of realizations) either for spontaneous or external stimulus-driven polarization out of 100 realizations.

1. **Co-alignment:** Polarity axes are roughly parallel to each other, with an angle difference less than 45 degrees.
2. **Collision:** Polarity axis of each cell points to the cell-cell junction; the axes are roughly antiparallel (parallel vectors with opposite directions) and point within a 36-degree angle opening about the horizontal line (orange sectional area in (2.) in Fig. 2d).
3. **Non-polarized:** Either one cell or both cells fail to polarize if either one of the networks never goes above threshold level *C*_crit_.
4. **Misalignment:** Neither one of the above cases, meaning that both cells do polarize, but their polarity axes are neither in co-alignment nor collision arrangement, as defined above.
5. **Supracellular** (or leader-follower) arrangement Previous work has reported before on the supracellular organization of motile groups of cells [56–59]. In our model, a supracellular arrangement is identified when the prospective leader’s polarity axis is aligned in any direction, but away from the cell-cell junction defined as a 45-degree contact region. Meanwhile, the prospective follower’s polarity axis is oriented toward the leader’s center-of-mass, within a 45-degree angle opening about the horizontal line. This configuration does overlap with co-alignment and misalignment outcomes.

In the absence of interactions between cells, there is approximately a 25% chance for the pair to co-align and thus, polarize in the same direction (Fig. 2e, Movie S1). Even if we account for intrinsic variability in the kinetic rates and parameters of either biochemical or mechanical signaling modules across cells, the co-alignment outcome is not improved (first seven rows in Table 1). The co-alignment outcome is also not significantly improved for signal-induced polarization – in this scenario, we consider that only one of the two cells receives an external stimulus which is locally enforced by a spatial profile for the binding/unbinding rates for Rac molecules along the plasma membrane (Fig. 2e). Similar findings hold for the supracellular arrangement. These results inform us that for coordinated symmetry breaking across a pair of cells, the cell-cell junction must mediate the front-to-rear polarity signaling module, but <u>which types of couplings</u> (inhibitory or excitatory) of <u>which signaling components</u> (Rho GTPases and/or actin network) <u>can improve co-alignment or supracellular alignment?</u>

### Speculated intercellular interactions for cell doublet polarity

We identify around a dozen mechanisms that have been proposed based on biological experiments (Table 1). However, in our model, we find that many of the speculated interaction models do not improve orientation in the same direction of the polarity axes of the doublet. We find that a symmetric, reciprocal interaction results predominantly in collision configuration. This is a likely outcome since, for example, increased Rac association at the cell-cell junction in both cells will lead to the formation of protrusive fronts at the cell-cell junction due to the positive feedback between Rac and branched F-actin. Symmetric reciprocal unbinding leads to similar results – for example, increased Rho unbinding, predisposes Rac binding at the cell-cell junction which will result in branched F-actin networks pointing towards each other at the intercellular region, and thus high likelihood of collision arrangements (Fig. 2d). We also consider concentration dependent interactions but find no significant differences in the outcomes. Our model findings for the literature speculated mechanisms are summarized in Table 1.

Due to the lack of positive results, we perform an exhaustive search of cell-cell couplings at the cell-cell junction that could favor co-alignment or supracellular arrangements. There are a large number of possibilities – for example, if we only consider down- or up-regulation of binding and unbinding kinetic rates (*k*_on_, *k*_off_ for both Rac and Rho) and network assembly rates (*k*_*A*_, *k*_*B*_) in the mechanical circuit that is 6 parameters with 5 choices (default, 10-fold decrease/increase, 1000-fold decrease/increase): 5^6^ = 15625 and then across two cells (15625*C*2) + 15625 = 122, 078, 125 combinations. The count does not include more complex schemes like concentration dependent changes or nonlinear interactions. Furthermore, every parameter choice requires multiple realizations due to the stochastic nature of the model. For computational feasibility, we proceed with an exploratory search of the simplest interaction pathways that involve the biochemical circuit only, the mechanical circuit only, or crosstalk between biochemical and mechanical circuits as identified in Table S2. For each entry in Table S2, a pathway of interaction is chosen – this includes interaction components, nature of interaction (inhibitory or excitatory), and strength – and 100 realizations for a particular interaction pathway are generated. The outcomes for each possible arrangement are listed in Table S2 and below, we only discuss the pathways that significantly improve co-alignment or supracellular arrangement of the polarity axes in the pair. We define a “favorable” interaction as an interaction that results in over 70% likeliness for either co-alignment or supracellular arrangement, as it represents roughly a three-fold increase over the co-alignment probability in the uncoupled case.

### Asymmetric crosstalk of the biochemical signaling circuit significantly improves doublet co-orientation of polarity axes

First, we ask whether collective orientation of polarity axes in the cell doublet can be established only through biochemical interactions of the small GTPases at the cell-cell junction. The biochemical interaction may involve direct molecular contacts between juxtaposed cells or indirect couplings mediated by diffusible molecules. Here, to enforce biochemical intercellular crosstalk, we assume either the binding or the unbinding rate of one GTPase is increased locally at the cell-cell junction in one or both cells (Fig. 3a). In the model, the implementation involves changing the kinetic rates in Eq. 1 but only locally at the cell-cell region. In Fig. 3a, we highlight which interactions lead to 70% or greater likeliness outcome for either co-alignment arrangement (1 in Fig. 3a) or supracellular arrangement (2 in Fig. 3a). We find only one type of interaction is favorable for co-alignment and supracellular arrangement – asymmetric regulation of Rho GTPases (Fig. 3b). Asymmetric regulation across the cell-cell junction could happen through one of 4 ways: binding (or unbinding) of one Rho GTPase in one cell and similar action of binding (or unbinding) of the complementary Rho GTPase in neighboring cell (first two boxes in Fig. 3b), or complementary kinetics, binding in one cell and unbinding in neighboring cell, of Rac (or Rho) (last two boxes in Fig. 3b).

**Fig 3.**
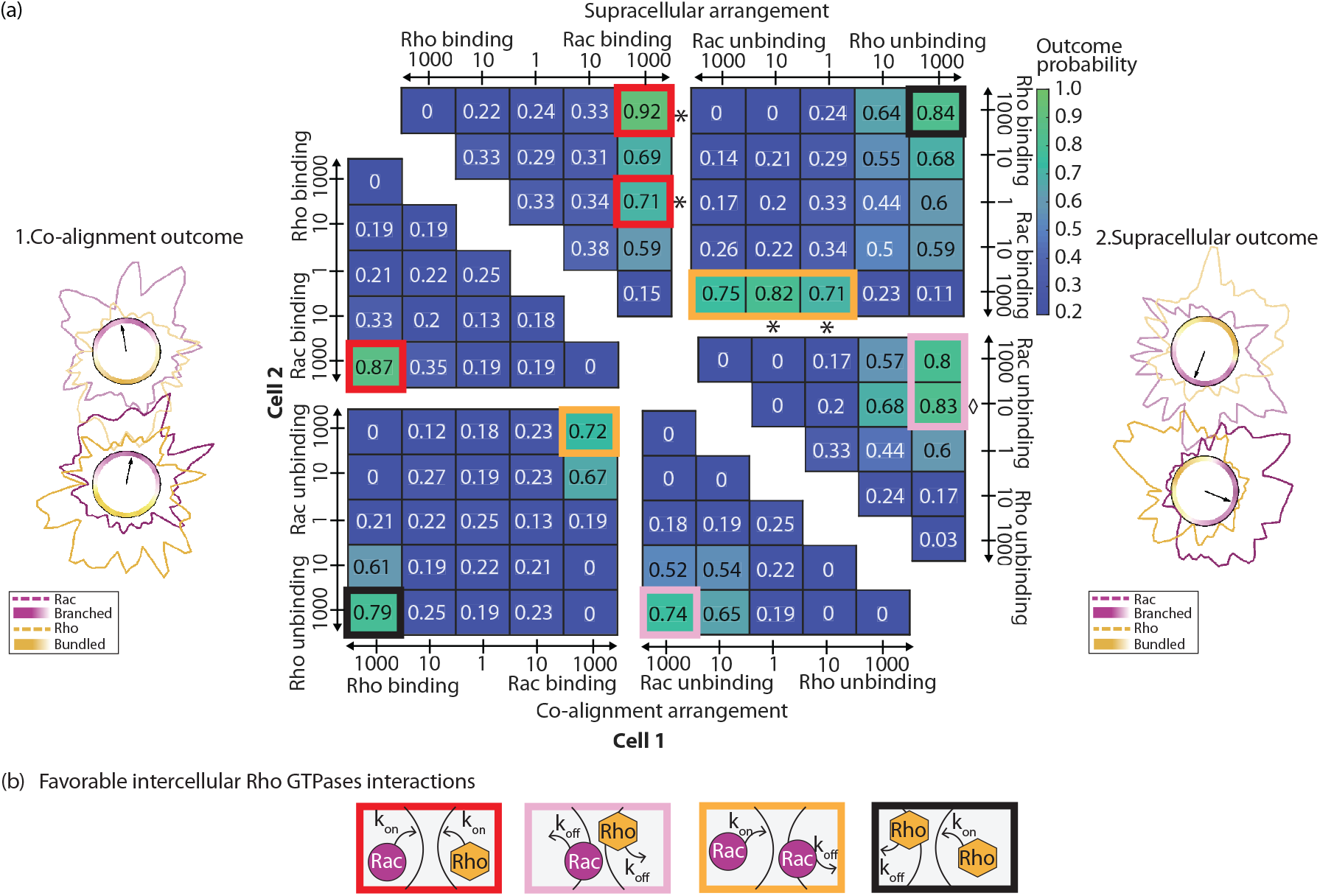
Only asymmetric regulation of Rho GTPases establishes co-oriented spontaneous polarization of the doublet. (a) Parameter sweeps for Rho GTPase interactions at the intercellular junction. All possible combinations of enhanced binding and/or unbinding of either Rac or Rho are considered. The outcome probabilities for either co-alignment (bottom) or supracellular (top) arrangement are presented for these interactions out of 100 realizations. The outlined boxes indicate 70% or greater probability for the arrangement. Sample simulation outputs, resulting in (1) co-alignment arrangement (Movie S2) and (2) supracellular arrangement (Movie S3). Black arrows show the front-to-rear polarity axes. (b) Emergent favorable Rho signaling pathways to achieve either co-alignment or supracellular arrangement in the absence of an external stimulus.

Irrespective of the type of asymmetric coupling, a probability of 70% or greater is attained for either co-alignment arrangement (bottom left, Fig. 3a) or supracellular arrangement (top right, Fig. 3a). For supracellular arrangement, we find that the region of favorable outcomes expands slightly. As an example, for supracellular arrangement, it is sufficient for Rac binding to be up-regulated in one cell only (* in Fig. 3a), while the conditions for the neighboring cell – either up-regulated Rho binding or Rac unbinding – can be relaxed. A similar statement cannot be made for Rho unbinding; if Rho unbinding is up-regulated in one cell, only Rac unbinding in the neighboring cell yields high likeliness of supracellular arrangement (⋄ in Fig 3a).

The Rho GTPases crosstalk could be mediated in a concentration independent or dependent way. Above, we review concentration independent interactions. Next, we explore whether collective polarization of doublets could be improved – either higher likeliness of co-alignment or supracellular outcomes or more pathways – by regulating GTPase kinetics proportional to the concentration of the molecule adjusted in the neighboring cell at the cell-cell junction. The concentration dependent interaction was implemented by multiplying the appropriate kinetic rate by the total amount of molecules in the neighboring cell engaged in that specific interaction pathway. For example, if the original intercellular interaction was enhanced binding of complementary Rho GTPases with 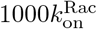 in cell 1 and 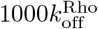 in cell 2, the concentration dependent version would be 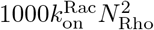 in cell 1 and 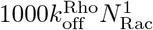 in cell 2. Here, 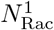 is the total number of active, membrane-bound Rac molecules, while 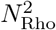 is the total number of active, membrane-bound Rho molecules in a small neighborhood around the neighboring point along the cell membrane in cell 1 and 2, respectively. Surprisingly, we find a significant drop in the likeliness of co-alignment of the doublets (bottom right, Fig. S1). This is because up-regulation of binding will be minimal if the corresponding molecule concentration is zero or low. In the case of supracellular arrangement, the results are qualitatively the same but notably asymmetric kinetics for Rac does not yield a probability outcome of 70% or greater. We conclude that in our model, concentration independent interactions are more likely to yield collective polarization in the doublet.

### Supracellular arrangement can be achieved through F-actin network regulation but co-alignment is less likely to be achieved

After considering intercellular communication of GTPase circuits, we ask whether co-orientation of polarity axes can be established only through interactions of neighboring F-actin networks at the intercellular region (Fig. 4). The dynamics of the polarity mechanism are initially described in Model section, except at the cell-cell junction, where the growth rates of F-actin networks are either up-regulated or down-regulated, independent or dependent on the concentration of actin networks in the neighboring cell. To enforce this crosstalk of F-actin networks, the equations for the mechanical circuit for cell *i* are modified to:

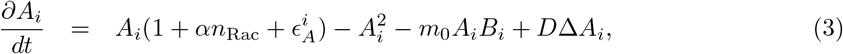

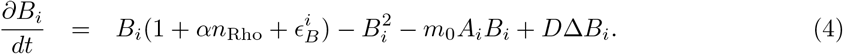

The subscripts *i, j* indicate the cell index (1 or 2 in a doublet). The newly introduced rates for the intercellular crosstalk between actin networks are 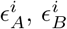 which can be either constant (Fig. 4a) or dependent on the local concentration of F-actin networks in neighboring cell *j* (Fig. 4d):

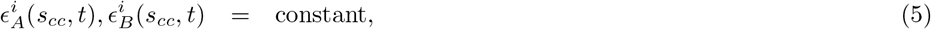

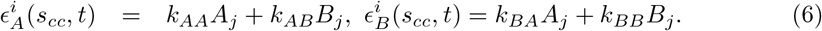

These newly introduced F-actin crosstalk rates are nonzero at the intercellular region, *s*_cc_, and zero everywhere else. All types of network couplings, including both diminishing (negative rate) and increasing (positive rate) are explored, and the results are summarized in Fig. 4b,c and Fig. 4e,f for concentration independent and concentration dependent rates, respectively. There are a total of 162 pathways: 162 = 3^4^ (3 choices: promote, inhibit, none; 4 parameters: 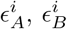 for *i* = 1, 2) + 3^4^ (3 choices: promote, inhibit, none; 4 parameters: *k*_*AA*_, *k*_*AB*_, *k*_*BA*_, *k*_*BB*_). The simulation results are shown for both co-alignment (Fig. 4b,e) and supracellular (Fig. 4c,f) arrangements, and the insets show probability values for selected outcomes. Fig. 4g shows the interaction schematic for the concentration dependent crosstalk that favors supracellular arrangement, with likeliness of 70% or larger.

**Fig 4.**
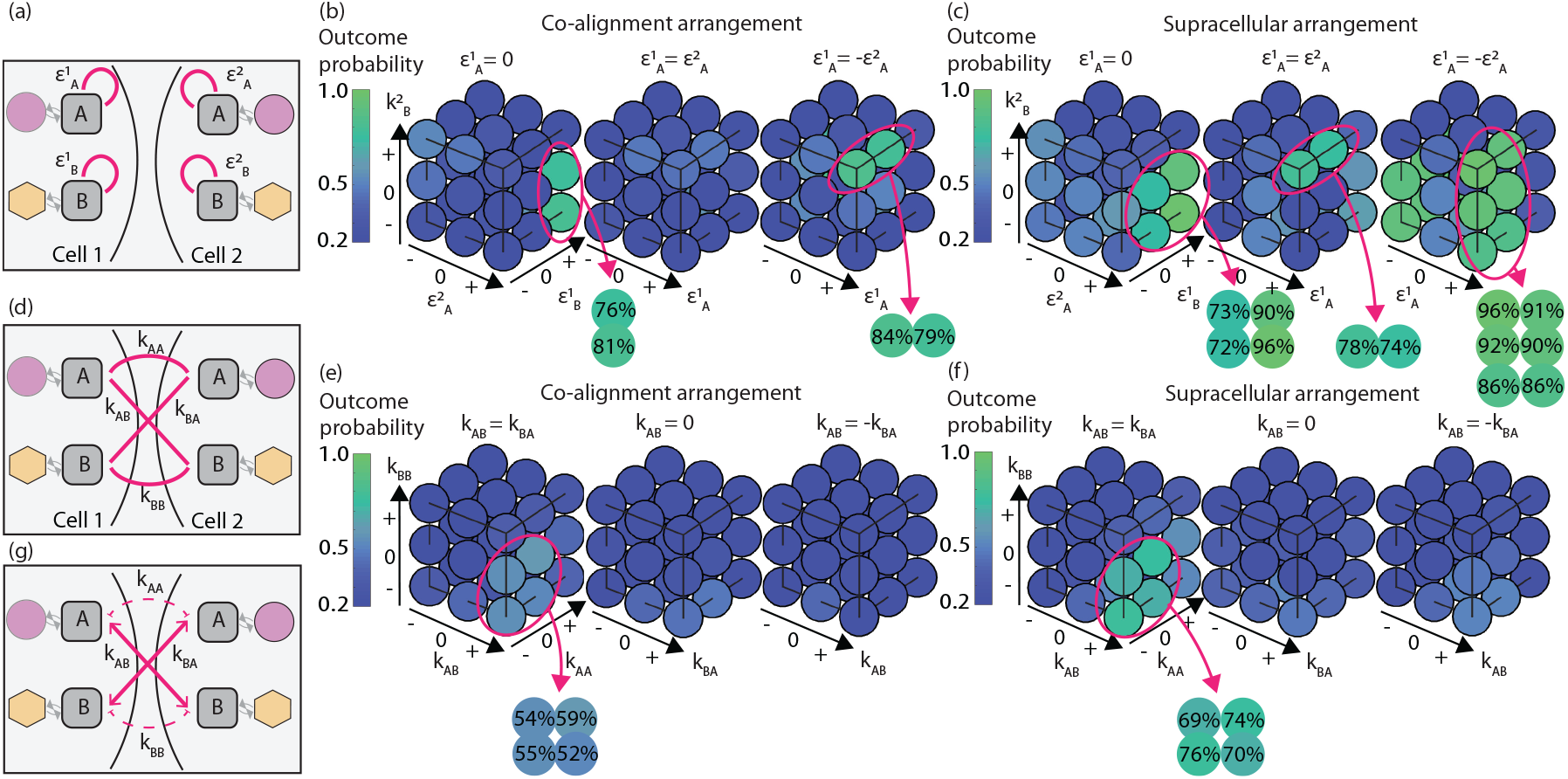
Crosstalk of F-actin networks gives rise to supracellular arrangement for doublet spontaneous polarization. (a) Schematic for concentration independent increase of network growth rates. Probability of (b) co-alignment arrangement and (c) supracellular arrangement for all possible combinations for an additive network growth rate constant which can be either positive, zero, or negative as in Eq. 5. (d) Schematic for concentration dependent increase of network growth rates. Probability of (e) co-alignment arrangement and (f) supracellular arrangement for all possible combinations for a concentration dependent network growth rate, as in Eq. 6. (g) Schematic of asymmetric network crosstalk interaction which solely favors only supracellular arrangement in the doublet.

We report on the model outcome in the case where network crosstalk rates are up-regulated in a concentration independent manner. We exhaustively consider all combinations of additive constants 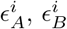which can take on either a positive (7.5), negative (-7.5), or zero value (Fig. 4b,c). The value of 7.5 is chosen as it represents 3/4 of the steady state concentration of either branched or bundled network concentrations. Again, the model does not support polarization of the doublet in the same direction with symmetric up-regulation of growth of branched or bundled networks at the cell-cell junction. The result holds for both co-alignment and supracellular arrangement. Co-alignment arrangement is most likely achieved with dual excitation-inhibition, meaning that growth of one network type is up-regulated in one cell while its growth is down-regulated in the second cell 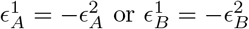. The likeliness of co-alignment outcome for these interaction pathways ranges between 81-84%. One side of this interaction pathway can be relaxed, to no regulation, and we find that the probability to establish co-alignment arrangement decreases slightly to 76-79% (Fig. 4b). Including any symmetric regulation collapses these favorable interactions; for example, 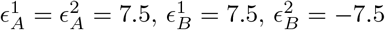 yields a 17% probability for co-alignment arrangement. Not surprisingly, the results are qualitatively similar for supracellular arrangement but with fewer constraints (Fig. 4c); for 70% or larger outcome likeliness it is sufficient to have only one network type engage in dual excitation-inhibition, while the other network type can engage in either none or symmetric interaction.

Results are quite different for concentration dependent interactions (Fig. 4d). A much smaller set of conditions allow for co-orientation of polarity axes in doublets and additionally, co-alignment outcome likeliness never reaches 70% (Fig. 4e,f). The model predicts that the probability outcome is maximized for the case of reciprocal (*k*_*AB*_ = *k*_*BA*_) and excitatory (*k*_*AB*_, *k*_*BA*_ *>* 0) couplings between branched and bundled networks in neighboring cells (Fig. 4e,f). Moreover, these favorable interactions require like-networks either engage in no interaction or inhibition (*k*_*AA*_, *k*_*BB*_ ≤ 0). In this scenario, the co-alignment arrangement is achieved in 52-59% of the cases (inset, Fig. 4e), but the likeliness of supracellular arrangement is higher, around 69-76%, with 0% cases of collisions (inset, Fig. 4f). The theme of our findings from crosstalk of Rho signaling circuits continues – co-orientation can be achieved with regulation of the formation of complementary networks across the cell-cell junction, a “push-n-pull” mechanism (Fig. 4g).

### Cellular variability does not change the set of intercellular interactions favors co-polarization

Can variability in the polarity machinery between cells in the doublet, either externally induced or intrinsically generated, could account for other regulatory mechanisms identified for co-orientation of polarity axes for collective polarization? Specifically, would intercellular conditions in either biochemical or F-actin network regulation change when cellular variability is considered? To test this hypothesis, we assume that one cell in the doublet has more responsive GTP activity though elevated baseline affinity in Rac and/or Rho association (Table S3) or greater baseline growth rate for contractile and/or protrusive network (Table S4). Given this asymmetry in the doublet, we scan a subset of the possible interactions at the intercellular junction and quantify the polarization outcomes. The subset of cell-cell regulatory interactions are: the 4 asymmetric Rho GTPase interactions schematically illustrated in Fig. 3b, elevated Rho GTPase mutual antagonism, CIL/COA, and F-actin network crosstalk (as in Fig. 4g). In summary, we find that cell-to-cell variability does not produce additional intercellular regulation mechanisms to ensure co-polarization of the doublet.

In the case of more responsive GTPase activity, we assume that one cell in the doublet has higher binding rate for either Rac or Rho or both GTPases – the rate is increased by a factor of 10 along the entire domain of the cell membrane before additional assumptions for intercellular communication are made. With small differences, co-alignment or supracellular arrangements are favored if the intercellular interaction is one of the 4 asymmetric Rho GTPase crosstalk ways or F-actin network crosstalk. Next, we consider cellular variability with respect to the F-actin dynamics – in one cell, we assume a higher network growth rate for either bundled, branched, or both actin networks:

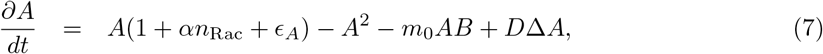

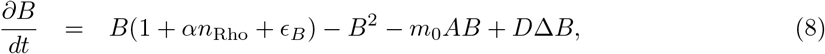

where either *ϵ*_*A*_ = 1, *ϵ*_*B*_ = 1, or both. The model results for mechanical variability are nearly the same as in the case of biochemical cellular variability; co-alignment or supracellular arrangements are favorable outcomes if the intercellular interaction is one of the 4 asymmetric Rho GTPase crosstalk or F-actin network crosstalk (Tables S3 and S4). There is one notable exception – if one cell has higher baseline growth rate of bundled networks, most of the asymmetric Rho GTPase or F-actin crosstalk ways do not lead to high probability of co-orientation of the polarity axes. In this case, the only favorable intercellular communication involves asymmetric Rho kinetics. Thus, we find a difference in intercellular communication pathways for collective polarization if there is cellular variability in the branched versus bundled network growth rates.

### The same set of intercellular interactions are favored for external stimulus-driven polarization in the doublet

To determine whether the doublet model exhibits sensitivity and adaptation to external signals, we simulated polarization in the presence of a directional bias. Trivially, in our model, if both cells receive the same external signal, any intercellular coupling, including no coupling, results in both cells polarized in the direction of the signal [35]. Instead, an external stimulus is available to only one cell, and we ask what type of cell-cell interactions, biochemical and/or mechanical, are required for co-alignment and/or supracellular configuration of polarity axes. To impose an external stimulus in one cell, we assume that the binding rates for Rac/Rho molecules are non-constant along the plasma membrane, which is equivalent to a directional bias, as shown in Fig. 5a – the Rac binding rate varies oppositely to the Rho binding rate, as the spatial complement of the curve: the sum of Rac and Rho binding rates is held fixed. The cell subject to an external stimulus is labeled as “cell 2”. As in [35], we report that in cell 2 a polarized state evolves from random initial conditions, with a Rac peak with the same orientation as the external bias (Fig. 2e), but not necessarily in the neighboring cell. An exhaustive search of intercellular pathways is performed and the probability for co-alignment and supracellular arrangement is reported for the doublet in the presence of an external stimulus applied to cell 2 (Fig. 5b-e). Briefly, we find qualitative differences between spontaneous and stimulus-induced co-polarization of the cell doublet. One important difference is that the pathway, based on F-actin network crosstalk as in Fig. 4g, is unlikely to yield either co-alignment or supracellular arrangement of polarity axes in the doublet.

**Fig 5.**
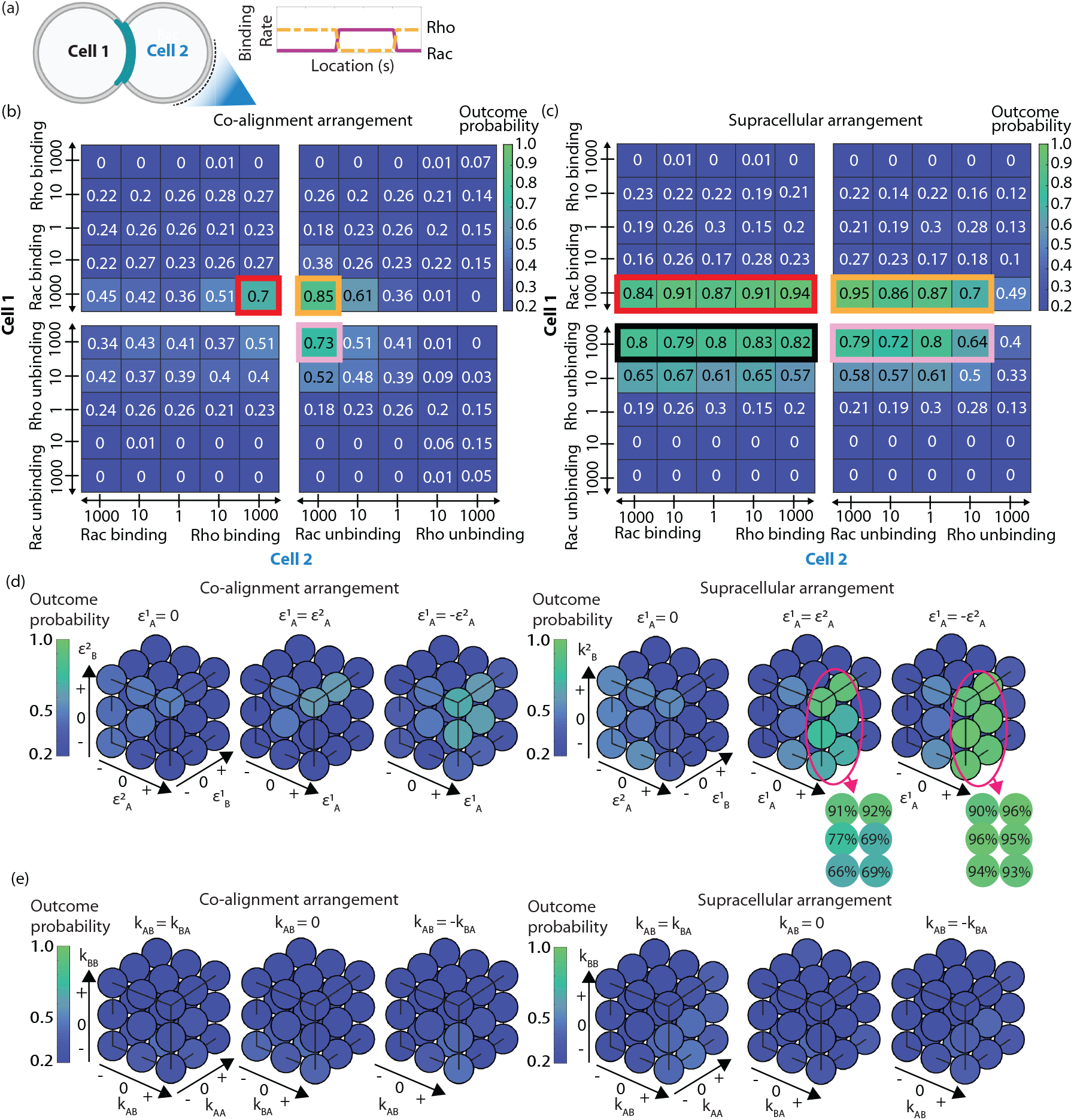
Results for signal-induced polarity in the doublet. (a) An external stimulus imposes a directional bias on the kinetic rates of both polarity proteins Rac and Rho. Rac binding (solid, magenta) and Rho binding (dashed, yellow) rates have the profile around the cell edge shown in this plot. (b-c) Probabilities for co-alignment and supracellular arrangement of the polarity axes of the doublet in the presence of an external stimulus, applied to cell 2, with intercellular couplings of Rho GTPases. The outlined boxes indicate 70% or larger likeliness for the arrangement. The color of the box outline matches the cell-cell interaction schematic in Fig. 3b. (d) Probability of co-alignment arrangement and supracellular arrangement for all possible combinations of additive network growth constants, which can be either positive, zero, or negative. (e) Probability of co-alignment and supracellular arrangement for all possible combinations of network-dependent growth rates.

#### Asymmetric regulation of Rho GTPases

In addition to the assumptions for GTPase kinetic rates in cell 2 due to the external stimulus, we enforce that neighboring cells engage in biochemical intercellular crosstalk by up-regulating either binding or unbinding rates of one GTPase in the cell-cell contact region (Fig. 5b,c, Movie S4). Unlike the spontaneous case, there are more cases to be explored, since the symmetry of the doublet is lost due to the fact that cell 2 is subjected to an external stimulus. An outline is drawn around the favorable interactions, defined as interactions which lead to 70% or larger likeliness of the outcome. The color of the outline indicates the mechanism for asymmetric Rho GTPase regulation, as illustrated in Fig. 3b.

For co-alignment arrangement (Fig. 5b), there are only 3, rather than 4, favorable interactions: (1) up-regulated binding rates of complementary GTPases, (2) complementary elevated kinetic rates (binding/unbinding) of Rac, or (3) up-regulated unbinding rates of complementary GTPases. Notably, the absent pathway is complementary elevated kinetic rates (binding/unbinding) of Rho. This case results in only ∼50% likeliness of co-alignment configuration; reduced since this would cause two competing fronts for cell 2 (or the absence of a rear). On the other hand, for supracellular arrangement (Fig. 5c), the favorable interactions are exactly the same as those identified for spontaneous polarization. However, there are even fewer constraints on cell 2 in these favorable interactions, as demonstrated by the region covered by outlined boxes. The only constraint is that in cell 2 the unbinding rate of Rho cannot be too high. In summary, the biochemical-based intercellular pathways in signal-driven polarization of the doublet approximately fall under the same umbrella of interactions identified for spontaneous polarization.

#### Push-n-pull of F-actin networks

In the case of F-actin network interactions, we report on the partial collapse of an intercellular interaction that was favored for spontaneous polarization. To demonstrate this result, in addition to the external stimulus assumption for a spatial profile for GTPase kinetic rates of cell 2, the doublet cells engage in F-actin crosstalk by inhibition or excitation of network growth rates in a concentration independent or dependent way at the cell-cell contact region. An exhaustive scan is done for all possible combinations of F-actin interactions and model outcomes are reported (Fig. 5d,e).

The highest probability for either co-alignment or supracellular arrangement is achieved with concentration independent altered network growth rates (Fig. 5d). Specifically, only the supracellular arrangement can be achieved with a probability of at least 70% – this occurs if in cell 1 at the cell-cell junction, formation of branched F-actin network is up-regulated 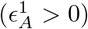, while the growth of bundled F-actin network is either not changed or down-regulated 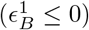 (Fig. 5d). This suggests that for supracellular arrangement, in the presence of an external stimulus, the cell not subject to a signal must up-regulate formation of the branched network at the cell-cell junction.

For concentration dependent assembly rates, we report that none of the considered interactions improve the probability past 50% for either co-alignment or supracellular arrangement of polarity axes of the doublet (Fig. 5e). We understand this result because of the dynamics of the F-actin network formation, which require a strong spatial constraint so that one network does not take over the entire domain. In our model, within an individual cell, the spatial constraint is provided by the segregation of the Rho GTPases. To see if the likeliness of co-polarization of the doublet can be improved, we include additional crosstalk between GTPases and F-actin networks by requiring that the association kinetic rates dependent on the local F-actin concentration in the neighboring cell. We consider only a subset of the interaction pathways and introduce GTPase-actin crosstalk – Rho binding rate depends on the concentration of branched network in the neighboring cell, and similarly, Rac binding rate depends on the concentration of bundled network in the neighboring cell. Surprisingly, the results are only marginally improved by including GTPases in the F-actin network crosstalk (Table S5).

### Response to switch in the orientation of the external stimulus

Lastly, we assess whether a polarized doublet shows sensitivity to a new signal direction by re-polarizing in the new orientation. We initiate the cell doublet by applying an external stimulus located in the lower right corner of cell 2, centered at 225^°^ degrees from the vertical. After a period of time (5 seconds), we remove the stimulus and place a new stimulus in the upper left corner of cell 1, centered at 45^°^from the vertical. Only a small subset of couplings at the cell-cell region are explored: (1) uncoupled, (2) up-regulated binding of complementary Rho GTPases, (3) asymmetric regulation of kinetic rates (binding/unbinding) of Rac, (4) concentration dependent up-regulated unbinding of complementary Rho GTPases, and (5) F-actin network crosstalk. The outcome probabilities for each of these interactions are summarized in Table S6. Regardless of the intercellular interaction, in most trials we find that after the signal switch, all polarity molecules dissociate from the plasma membrane (Movie S5). This motivates a second implementation of the signal switch; if all Rac and/or Rho molecules unbind, a neutralization process is initiated. The neutralization process assumes re-initialization of GTPases by resetting 10% of Rac/Rho molecules active and bound to the plasma membrane in random locations. Within the subset of interactions tested, we find high probability of either co-alignment or supracellular arrangement for concentration dependent of up-regulated unbinding of complementary Rho GTPases at the cell-cell junction (Table S6). In this pathway, in the cell subjected to the external stimulus, Rac unbinding rate is elevated while Rho unbinding rate is elevated in the other cell, depending on the Rho and Rac concentrations, respectively, in the opposite cell. The scenario produces 55% of pairs co-aligning to the signal and 84% displaying a supracellular arrangement (Movie S6). This is the same pathway that gave favorable results in the first implementation of the signal switch.

### Time for a co-polarized state in the doublet is not reduced compared to an individual cell

To determine whether a doublet can break symmetry more readily than an individual cell, we quantify the time to reach a polarized state for single cells and doublets with various intercellular interactions. All 5 favorable intercellular interactions are considered, which include the 4 cases of asymmetric Rho GTPase regulation (Fig. 3b), and the F-actin network crosstalk (Fig. 4g). The time to reach a polarized state either within a cell or as a cell doublet is reported in Fig. 6. In our model, we find that the doublet always takes longer or just as long to break symmetry when compared to a single cell.

**Fig 6.**
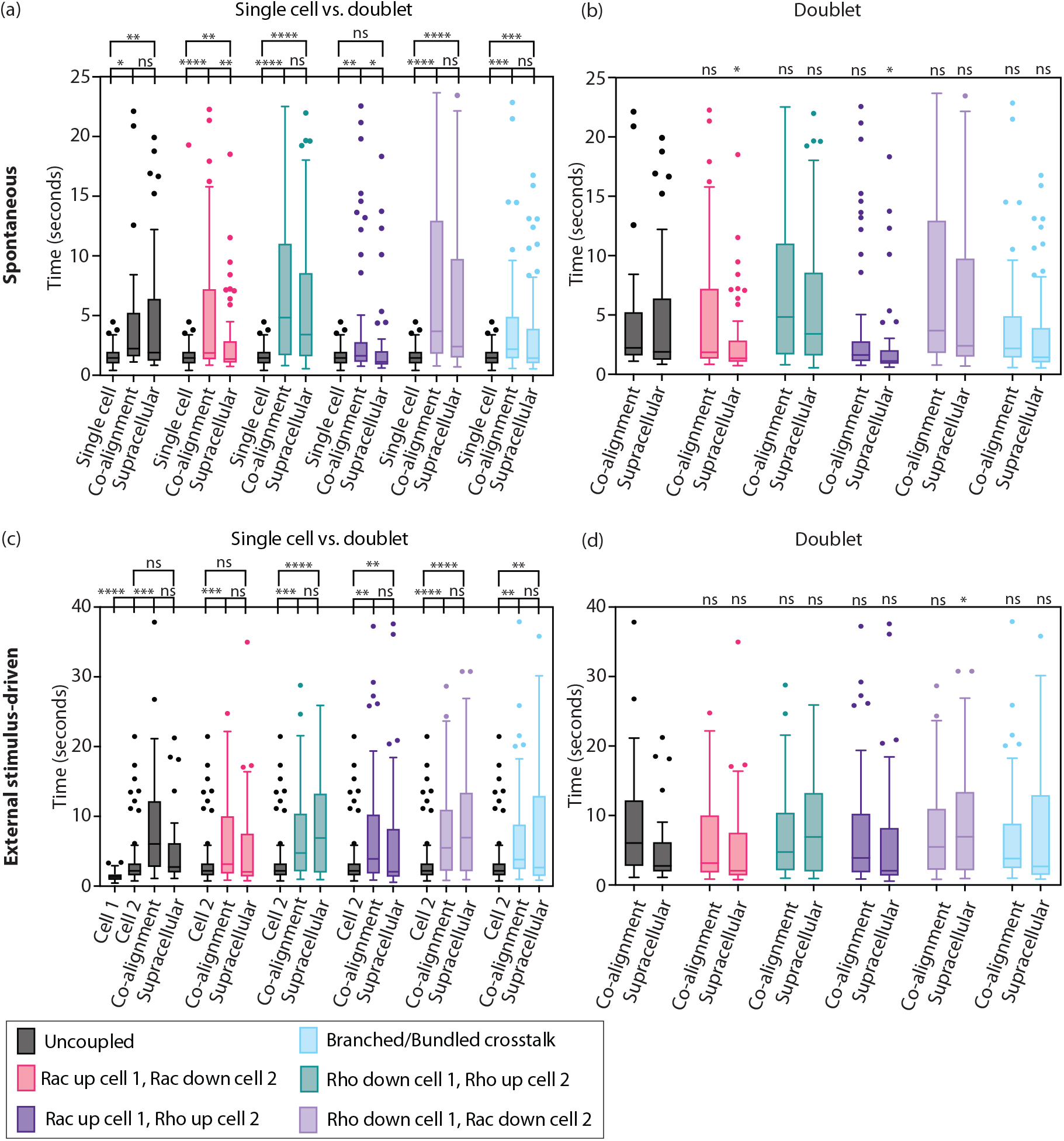
Time to co-polarize for a doublet is not reduced compared to that for an individual cell. (a) Comparison of time to reach a polarized state in the absence of any external stimulus of single cell against doublet in either co-alignment or supracellular organization. (b) Comparison of time to reach polarized state in the absence of any external stimulus of uncoupled doublet against doublet with an intercellular interaction. (c) Comparison of time to reach polarized state with an external stimulus given to cell 2 against doublet in either co-alignment or supracellular organization. (d) Comparison of time to reach polarized state in the presence of an external stimulus of uncoupled doublet against doublet with an intercellular interaction. Each color represents a particular interaction at the cell-cell junction region. The number of cases considered per interaction is 100. Welch’s ANOVA; n.s. p≥ 0.05, ^*^p< 0.05, ^**^p< 0.01, ^***^p< 0.001, ^****^p< 0.0001. Boxes are the 25th to 75th percentiles, and bars indicate ± interquartile range.

A polarized cell state is defined as in Model section – in 100 realizations with random initializations, a polarized cell state is achieved when both branched and bundled concentrations are above the threshold level, and the orientation of the polarity axis is from the cell centroid to the middle point of the threshold region for branched F-actin network. Furthermore, for reporting time to reach a polarized state, we ensure that the orientation of the axis remains constant. This is especially relevant for doublets where the relative orientation of the polarity axes is important. Time to reach a fixed orientation of the polarity axis is the first instance when, within 100 time steps, the consecutive angle difference of the axis does not change more than a small amount of *π/*12 radians. For individual cells, we find that the time for polarized state is longer with an external stimulus than without (spontaneous) (uncoupled in Fig. 6a, gray). We attribute this outcome to the loss of bidirectional feedback between Rho GTPases and F-actin networks since in the external stimulus scenario the GTPases are no longer dependent on the F-actin network concentrations but rather have spatially fixed rates.

For doublets, all 5 cell-cell couplings are evaluated in the absence (Fig. 6a,b) and presence (Fig. 6c,d) of an external stimulus. Three sets of comparisons are performed; Figs. 6a and c show comparisons of time to polarize of individual cell against doublet and also time to polarize doublet in co-alignment against supracellular arrangement. Figs. 6b and d show the comparison of the time to polarize doublet with no coupling against one of the 5 cell-cell couplings. First, we find that the time to reach a polarized state is as long or longer than an individual cell, indifferent of the presence or absence of an external stimulus. Further, that is true, indifferent of whether we look for co-alignment or supracellular arrangement of polarity axes. Two couplings stand out as situations for which there is no statistically significant difference in the polarization time: asymmetric enhanced binding of complementary Rho GTPases (Fig. 6a, dark blue) for supracellular arrangement with no external signal and asymmetric elevated binding/unbinding of Rac (Fig. 6b, red) also for supracellular arrangement but with signal-induced polarization. Second, for a few cell-cell couplings, it is faster to achieve a polarized state in supracellular arrangement over co-alignment arrangement (Fig. 6a, red and dark blue). Surprisingly, that is the case only for spontaneous polarization; in the case of stimulus-driven polarization, there is no statistically significant difference in polarization time between the two arrangements. Third, we find that that intercellular couplings not only ensures higher likeliness of co-orientation of the cell group in the same direction but also reduce the time to achieve a polarized state over the uncoupled scenario (Fig. 6b and d). For spontaneous polarization, the time to achieve supracellular arrangement can be reduced through two intercellular interaction pathways – either asymmetric regulation of Rac kinetics (Fig. 6b, red) or enhanced binding of complementary Rho GTPases (Fig. 6b, dark blue). With an external stimulus, supracellular arrangement is achieved faster through enhanced binding of complementary Rho GTPases (Fig. 6d, violet).

### Geometric arrangement affects organization of polarity axes in larger groups

Finally, we report our findings for mechanisms for co-orientation of polarity axes for groups of 4 cells. As we did above, all 5 favorable intercellular interactions are considered, which include 4 cases of asymmetric Rho GTPase regulation and the F-actin network crosstalk. We find that the model results regarding the success of co-orientation also depends on the group’s initial geometric arrangement (Fig. 7).

**Fig 7.**
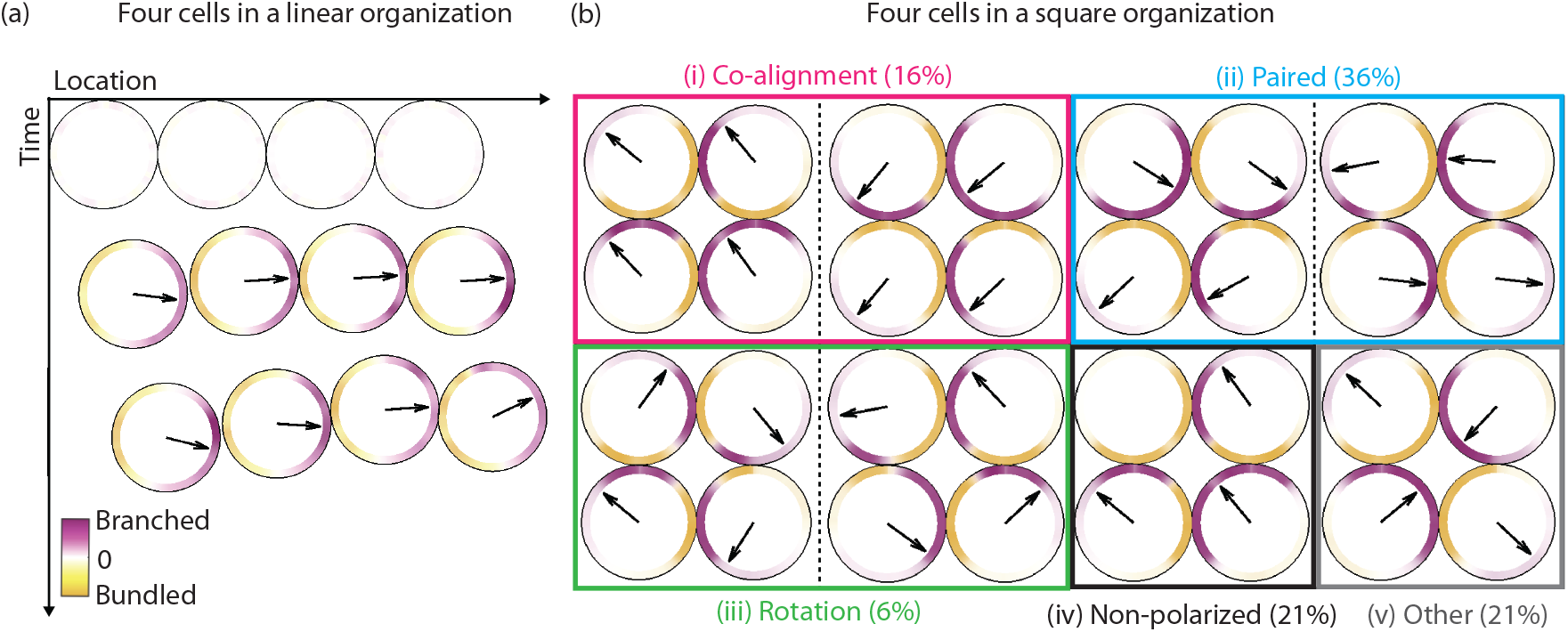
Polarization outcomes for 4 cells placed initially in two different geometric arrangements. (a) Time evolution of a 4-cell cluster in a linear arrangement, where each cell in the cluster moves with a constant velocity in the direction of its polarity axis. (b) Possible outcomes of a 4-cell cluster in square arrangement. The probabilities are computed from 100 realizations of the quadruplet in a square configuration with F-actin network crosstalk implemented at all cell-cell junctions.

When cells are placed in a linear arrangement (Fig. 7a, Movie S7), we find very similar outcomes compared to cell doublets – asymmetric regulation of Rho GTPases is favored as it results in high probability of co-alignment or supracellular arrangements. Unlike cell doublets, the crosstalk of F-actin networks is not sufficient to produce either co-alignment or supracellular arrangement (Table S7). A second initial geometric arrangement of the quadruplet considered is a square arrangement (Fig. 7b). In the absence of cell-cell couplings, a wider range of orientation of polarity axes is permissible due to a seemingly larger degree of freedom in the configuration. As an example, we categorized the outcomes of 100 realizations of a quadruplet in square arrangement with F-actin crosstalk at each cell-cell junctions (Table S8). The crosstalk of F-actin networks is implemented as the local formation of branched F-actin network in one cell is enhanced by the local presence of bundled F-actin network in the neighboring cell (Eqs. 3-4 and 6). A scan of the simulations reveals that there are 5 possible configurations of polarity axes in the quadruplet: co-alignment, paired alignment, circular (clockwise or counterclockwise) alignment (Movie S8), misalignment, or non-polarized. Overall, we find that co-alignment is rarely achieved. Of the 100 realizations, the 5 possible configuration of polarity axes are distributed as follows: 16% co-aligned, 36% paired, 6% circular, 21% misaligned, and 21% non-polarized in at least one cell (Fig. 7b).

## Discussion

The initiation of collective cell migration involves a complex web of signaling pathways and cytoskeletal rearrangement. In this Article, in one particular cell polarization model, we find that only asymmetric intercellular regulation of Rho signaling, or cytoskeletal dynamics can give rise to congruent orientation of polarity axes of cells in a doublet (Fig. 8). We come to this conclusion by examining all possible (linear) interactions at the cell-cell junction of either kinetic rates of Rho GTPases and/or F-actin network growth. The general question of how symmetric junctional proteins, like cadherins, establish asymmetric regulation remains a rich and active area of research. Within this framework the nature of the coupling, direct or indirect, is abstracted away, and instead we think of its downstream effect on the Rho GTPase signaling pathway and/or formation of F-actin networks locally at the cell-cell junction [29].

**Fig 8.**
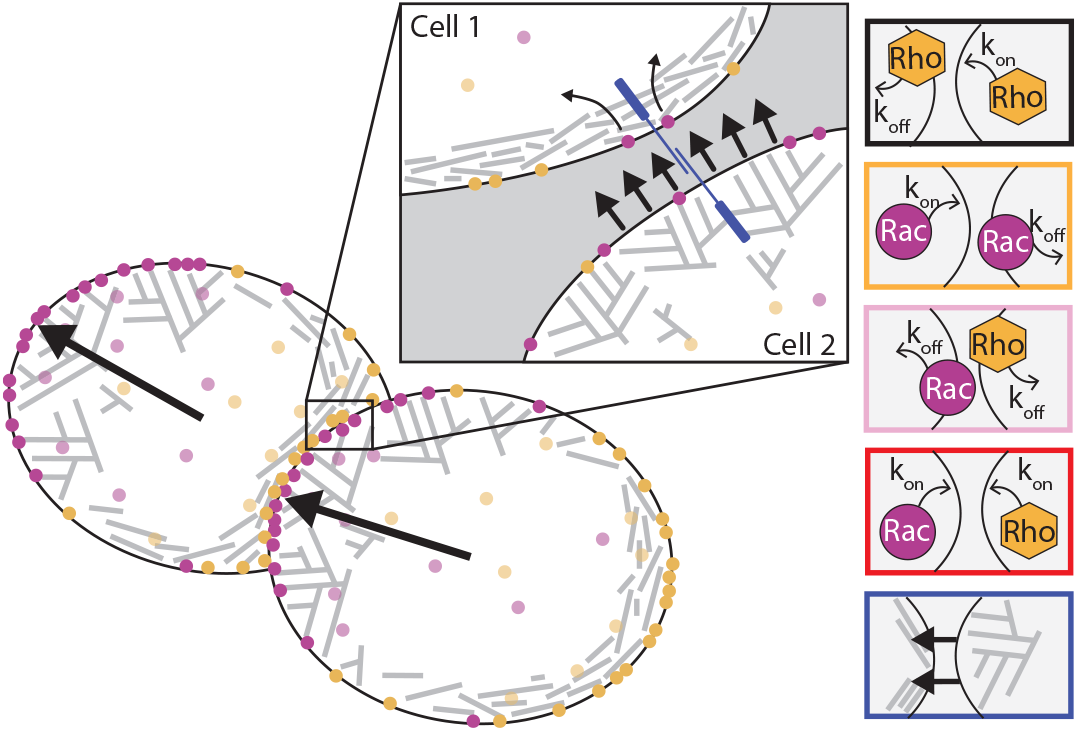
Illustration of the working principles underlying the set of intercellular interactions for collective orientation of polarity in a pair of cells. Inset illustrates our hypothesis that protrusive forces could drive enhanced dissociation of Rac in neighboring cell through mechanosensitive junction proteins.

While in certain cellular systems polarity can arise from only signaling or mechanical pathways, many cells rely on the interplay between the two to robustly break symmetry to initiate locomotion. Furthermore, we are motivated to explore whether polarization is coordinated through F-actin and/or signaling dynamics in groups of cells. Therefore, we employ a mechanochemical model as the simplest polarization model where feedback between the biochemical and mechanical circuits gives rise to symmetry breaking without additional mechanical effects [60, 61] or unverified kinetic details [62]. The biochemical circuit has three important features: active-inactive cycling of GTPases, difference in diffusion coefficients between plasma membrane and cytosol, and conservation of mass. The model results in the emergence of multiple peaks of activity without global cellular polarization; different types of molecules segregate locally, but the clusters of the two kinds do not aggregate in the respective halves of the cell, as required for establishment of a polarity axis for migration. This suggests to us that further feedback is needed. The second circuit is for two types of dynamic F-actin networks at the cell edge, a branched protruding meshwork and an actomyosin contractile bundled network. These networks spread slowly and randomly around the cell edge, due to physical movements and tread-milling of actin filaments, and turn over while maintaining a certain equilibrium density. The nontrivial interaction between these networks is competition, such that the local density of one tends to diminish the density of another. This interaction stems both from mechanical effects and from competition for molecular resources. It was shown in [13] that this competition between two actin networks is an important part of the spontaneous polarization process, but without cell movement, the model is not able to polarize the cell, as one network will always win. Positive, bidirectional, local, and linear feedback between the mechanical and biochemical circuits is sufficient for spontaneous polarity [35].

Since without any signal from the environment, the orientation of the front-to-rear axis is not pre-determined, we use the model to probe the intercellular interactions for the front-to-rear axes in cell doublets to point in the same direction – co-polarization. Out of the over 300 distinct pathways we scan, we find that only one type of interaction produces high probability of co-polarization of the doublet. This pathway involves asymmetric regulation across the cell-cell junction. The asymmetric regulation of Rho GTPases required to lead to spontaneous co-polarization can be achieved through one-sided dimming/suppression of the same type of Rho GTPase across the junction, which indirectly promotes the activation (and association) of the antagonistic Rho GTPase. Essentially, the mechanism can be re-stated in the terminology of the inhibition of “frontness” and “backness” implicated in polarization of neutrophils and *Dictyostelium discoideum*. At the cell junction, presence of “frontness” in one cell ensures diminished “frontness” in the neighboring cell and, thus, “backness” in the neighboring cell. Another mechanism is through enhanced activation of opposing Rho GTPases across the junction; for example, increased activation of Rac in one cell and enhanced Rho association in the neighboring cell results in co-polarization of the doublet. Co-polarization of the doublet can also be achieved with asymmetric regulation of mechanical circuits, namely up-regulation of growth of one type of network in one cell with simultaneous down-regulation of growth of the same network in the neighboring cell. We posit that this type of interaction could arise from displacement-induced behavior not dissimilar from what has been reported in keratocytes where the forward movement of the plasma membrane engages in positive feedback with assembly of the branched F-actin network in the lamellipodium [13]. This is not an opaque finding; however, the novelty of our work lies in the fact that at least in this theoretical framework, it is the only possible way to ensure cells in the group “agree” on the same axis of migration.

While the novelty of our work lies in the exhaustive search of possible intercellular interactions motivating collective polarity at the onset of migration, our model findings are aligned with recent experimental findings. In *Drosophila* border cells, leader-driven suppression of protrusive activity in follower cells is mediated by Rac [58]. In *Drosophila* follicular epithelial cells, Fat2 localizes to the trailing edge of each cell and promotes the formation of F-actin rich protrusions at the leading edge of the cell behind [40]. Similarly, human umbilical vein endothelial cells have been shown to have polarized Arp2/3 and VE-cadherin rich membrane protrusions, called “cadherin fingers”, which locally increase actomyosin contractility in follower cells as means for tissue level organization. Our model also supports that asymmetric regulation of Rho GTPases is a universal, albeit not exclusive, pathway to negotiate front-to-rear alignment across cell groups. Additionally, we find that co-polarization can also be achieved with crosstalk between the mechanical and biochemical circuits; either co-alignment or supracellular arrangement is a likely outcome if we assume that bundled network up-regulate Rac association in the neighboring cell and, similarly, branched network up-regulates Rho association rates (Table S2). The assumption that branched filaments through Arp2/3 can locally up-regulate Rac binding in the cell-cell region has been experimentally observed in epithelial cells [45]. In our model, we find that these asymmetric intercellular interactions are a conserved set of co-polarization pathways even with cellular variability or with external signal bias of the kinetics of the polarity molecules in a certain direction.

In addition to the parameter constrains of the polarization model for a single cell as described in [35], we find that the asymmetric regulation of the kinetic rates or network growth rates at the intercellular region has to be strong enough to overcome the other dynamics including feedback from F-actin networks but also GTPase active-inactive cycling. It is very likely that the model performs in 2D and 3D as well as in 1D, but neither the single cell model nor the doublet model has been extended to higher dimensions. Another limitation of our model is that more complex intercellular interactions are largely not considered – these could be nonlinear in nature or involve other mediators, such as curve-sensing proteins.

The first conceptual prediction of our model is that in the absence of the mechanosensitive response of intercellular junction proteins, co-orientation of polarity axes in the same direction is lost. For cell doublets that rely on cadherins for tissue organization, if their cadherins lacked cytoplasmic domains to engage with the cytoskeleton network, the doublets would result in poor co-orientation of their polarity axes with a 25% chance of co-alignment polarization arrangement. Interesting, one could conceive a setup where only one of the cells in the doublet has cytoplasmic domain negative cadherins. In this scenario, we posit that co-polarization would be successful only if the cadherin cytoplasmic domain negative cell in the doublet is exposed to an external stimulus, as in Fig. 6. This is because the neighboring cell still has mechanosensitive cadherins and can respond to the “backness” of the cell exposed to the external stimulus. Another conceptual biological prediction of our model is that the time for spontaneous polarization is not reduced for doublets over individual cells (Fig. 6). While studies have reported on the improved persistence of polarization in groups over individual cells [63, 64], here we report on the time to break symmetry in collective groups over individuals. Our model also posits that larger cell groups require additional or more complicated crosstalk to ensure co-polarization of the group, since our 4-cell cluster in a square arrangement could give rise to paired-like or rotational-like arrangements of polarization axes (Fig. 7b). Lastly, and not surprisingly, our model finds it is easier to achieve supracellular arrangement over co-alignment as this is more restrictive in terms of the orientations of polarity axes in the doublet.

We do not claim that our model can predict the biological details of co-polarization of groups of any cell type. Notably, one limitation of our model comes back to the underlying single cell polarization model: It is possible for our model to rely on other forms of feedback between the biochemical and mechanical circuits or even solely one of the two circuits. For example, negative, instead of positive, feedback between Rac and branched actin and Rho and actomyosin, respectively, could do the job [18, 48]. We also limited the dynamics of the model to the local chemical and mechanical processes, but global mechanical effects, for example, membrane tension, could play an important role in polarization of some cell types [60]. Another paradigm for mechanochemical polarization requires transport of chemicals in the signaling framework. The key to such models is that myosin-driven flow assists the polarization of signaling proteins by mechanically triggering the formation of a stable asymmetric chemical distribution [20, 65, 66]. Our model is simpler because it does not have directional movement – either in the form of a flow, as in these models, or in the form of whole cell movement, as in [13]. More detailed and complex models have included the cell-surface adhesion dynamics as a mechanical component in the biochemical polarization pathway [61].

Furthermore, the model does not include many molecular players – PIP, PI3K, PTEN, cadherins, G-proteins, actin regulators – but simply conceptually captures their lumped effect on the crosstalk between Rac/Rho and actin/actomyosin. Similarly, higher order, nonlinear interactions involving Hill-type functions are ignored. Instead, our model posits one of the simplest quantitative frameworks, avoiding additional assumptions, for understanding a possible mechanism for coordination of spontaneous polarization in a cell doublet prior to migration. We hope our model adds to the conversation on the effects of intercellular junction proteins on the polarity molecules and their downstream effectors.

## Supporting information

**S1 Video. Simulation of an uncoupled cell doublet with misalignment arrangement of the polarity axes**. The branched and bundled F-actin networks are plotted inside the cell, while the GTPase concentrations are shown outside the cell membrane (dashed). The Rho GTPase concentrations in cell 2 (right) are shown with transparency for visibility. The front-to-rear axis is shown in a black arrow. The time is shown in seconds.

**S2 Video. Simulation of a cell doublet which results in co-alignment arrangement**. The cell-cell coupling implemented is asymmetric up-regulated binding of complementary Rho GTPases, Rac in cell 1 and Rho in cell 2. The branched and bundled F-actin networks are plotted inside the cell, while the GTPase concentrations are shown outside the cell membrane (dashed). The Rho GTPase concentrations in cell 2 (right) are shown with transparency for visibility. The front-to-rear axis is drawn with a black arrow. The time is shown in seconds.

**S3 Video. Simulation of a cell doublet which results in supracellular arrangement**. The cell-cell coupling implemented is asymmetric up-regulated binding of Rac; Rac binding is up-regulated in cell 1 at the cell-cell junction, but nothing is changed in cell 2. The branched and bundled F-actin networks are plotted inside the cell, while the GTPase concentrations are shown outside the cell membrane (dashed). The Rho GTPase concentrations in cell 2 (right) are shown with transparency for visibility. The front-to-rear axis is drawn with a black arrow. The time is shown in seconds.

**S4 Video. Simulation of a cell doublet in the presence of an external stimulus**. Cell 2 is subjected to an external stimulus and the cell-cell coupling is asymmetric up-regulated binding of Rho GTPases. The resulting outcome is co-alignment of the doublet. The branched and bundled F-actin networks are plotted inside the cell, while the GTPase concentrations are shown outside the cell membrane (dashed). The Rho GTPase concentrations in cell 2 (right) are shown with transparency for visibility. The front-to-rear axis is drawn with a black arrow. The time is shown in seconds.

**S5 Video. Simulation of a cell doublet’s failed response to a signal switch**. Cell 2 is subjected to an external stimulus and the cell-cell coupling is asymmetric up-regulated binding of Rho GTPases. The resulting outcome is co-alignment of the doublet, as in Video . At time *t* = 5 seconds, the location stimulus is changed from cell 2 to cell 1 in the opposite direction. The doublet fails to re-polarize in the new direction. The branched and bundled F-actin networks are plotted inside the cell, while the GTPase concentrations are shown outside the cell membrane (dashed). The Rho GTPase concentrations in cell 2 (right) are shown with transparency for visibility. The front-to-rear axis is drawn with a black arrow. The time is shown in seconds.

**S6 Video. Simulation of doublet successfully re-polarizing in a new direction in response to a signal switch**. The setup is the same as video S5 but the after the signal switch, the doublet successfully re-polarizes in the new direction with co-alignment arrangement.

**S7 Video. Simulation of spontaneous polarization in co-alignment arrangement of 4 cells started in a linear configuration**. At cell-cell coupling, the interaction implemented is alternating asymmetric up-regulated binding of Rho GTPases. The branched and bundled F-actin networks are plotted inside the cell, while the GTPase concentrations are shown outside the cell membrane (dashed). The front-to-rear axis is drawn from the cell center to the median of the branched F-actin network above a threshold concentration (black arrow). Each cell domain moves with a fixed speed in the direction of the front-to-rear axis. The time is shown in seconds.

**S8 Video. Simulation of spontaneous polarization resulting in clockwise rotation of 4 cells in a square configuration**. The setup is the same as video S7, but the cells are started in a square configuration.

**A1 Appendix**. Model details and implementation. Figs. S1, S2, and Tables S1 to S8.

## Acknowledgments

We are grateful to Alex Mogilner and Andreas Buttenschön for helpful discussions and close reading of our manuscript. We thank Denise Montell and Yu-Li Wang for their insights into our model findings. We acknowledge the use of BioRender.com through Northeastern University access.

